# Informing virtual clinical trials of hepatocellular carcinoma with spatial multi-omics analysis of a human neoadjuvant immunotherapy clinical trial

**DOI:** 10.1101/2023.08.11.553000

**Authors:** Shuming Zhang, Atul Deshpande, Babita K. Verma, Hanwen Wang, Haoyang Mi, Long Yuan, Won Jin Ho, Elizabeth M. Jaffee, Qingfeng Zhu, Robert A. Anders, Mark Yarchoan, Luciane T. Kagohara, Elana J. Fertig, Aleksander S. Popel

## Abstract

Human clinical trials are important tools to advance novel systemic therapies improve treatment outcomes for cancer patients. The few durable treatment options have led to a critical need to advance new therapeutics in hepatocellular carcinoma (HCC). Recent human clinical trials have shown that new combination immunotherapeutic regimens provide unprecedented clinical response in a subset of patients. Computational methods that can simulate tumors from mathematical equations describing cellular and molecular interactions are emerging as promising tools to simulate the impact of therapy entirely *in silico*. To facilitate designing dosing regimen and identifying potential biomarkers, we developed a new computational model to track tumor progression at organ scale while reflecting the spatial heterogeneity in the tumor at tissue scale in HCC. This computational model is called a spatial quantitative systems pharmacology (spQSP) platform and it is also designed to simulate the effects of combination immunotherapy. We then validate the results from the spQSP system by leveraging real-world spatial multi-omics data from a neoadjuvant HCC clinical trial combining anti-PD-1 immunotherapy and a multitargeted tyrosine kinase inhibitor (TKI) cabozantinib. The model output is compared with spatial data from Imaging Mass Cytometry (IMC). Both IMC data and simulation results suggest closer proximity between CD8 T cell and macrophages among non-responders while the reverse trend was observed for responders. The analyses also imply wider dispersion of immune cells and less scattered cancer cells in responders’ samples. We also compared the model output with Visium spatial transcriptomics analyses of samples from post-treatment tumor resections in the original clinical trial. Both spatial transcriptomic data and simulation results identify the role of spatial patterns of tumor vasculature and TGFβ in tumor and immune cell interactions. To our knowledge, this is the first spatial tumor model for virtual clinical trials at a molecular scale that is grounded in high-throughput spatial multi-omics data from a human clinical trial.

## Introduction

### General information and clinical trial results for HCC

Worldwide, more than 900,000 people are diagnosed with liver cancer annually and more than 800,000 people die from it^1^. Hepatocellular carcinoma (HCC), the most common type of primary liver cancer, constitutes over 90% of all cases^2^. Over 70% of HCC tumors are unresectable at diagnosis stage due to local metastasis and limited hepatic function^3^. Even though only a small fraction of patients are eligible for hepatectomy or liver transplantation, they remain standard curative treatments for HCC. Recently, systemic treatments for HCC have been approved by the U.S. FDA. Immune checkpoint inhibitors (ICI), including nivolumab, atezolizumab, and pembrolizumab, target programmed cell death protein 1 (PD-1) or its ligand PD-L1 to promote anti-tumor immunity. Anti-angiogenic therapies, including regorafenib, cabozantinib, and ramucirumab, inhibit signaling of vascular endothelial growth factor receptor (VEGFR) and other angiogenic receptors, preventing neovascular formation in the tumor microenvironment (TME)^4^. To further improve treatment outcomes of systemic monotherapy in advanced stage HCC setting^5,6^, combination therapies are currently being examined for patients with HCC^4,7–9^. The pathological responses differ among patients and objective response rates range from 24% to 50%^10^. The pervasive heterogeneity in patient responses and numerous therapeutic agents being evaluated would require extensive combination clinical trials on large patient populations for comprehensive assessment of these new therapeutic strategies. New approaches are needed to distinguish the molecular and cellular states that discriminate responders and non-responders for personalized therapeutic selection at scale.

Computational models simulating tumors and their therapeutic response provide promising alternatives to address the limitations of human clinical trials. These model systems encode prior biological knowledge of how cells interact during tumor growth and in response to therapy into sets of equations. Solving these equations can then simulate the cells of a tumor over time, enabling comprehensive querying of the molecular and cellular states over the duration of treatment in a manner that is not feasible in humans or any current biological modeling framework. One powerful example of a computational model of tumors is Quantitative System Pharmacology (QSP) models, which mechanistically simulate disease progression processes, pharmacokinetics (PK), and pharmacodynamics (PD) of selected drugs. These models enable use of computational simulations for virtual clinical trials, and have become increasingly indispensable techniques for drug discovery and clinical trial design^11,12^. QSP models have been applied to analyze different types of cancer with various immune checkpoint inhibitors^11,13^. We have developed QSP platforms to investigate systemic therapies and anti-tumoral response at whole organ level for non-small cell lung cancer (NSCLC)^14^, breast cancer^15,16^, colorectal cancer^17^, and HCC^18^. However, due to a lack of spatial resolution, outputs from QSP models cannot be fully compared with quantitatively analyzed histopathological samples from tumors, including measures of intratumoral heterogeneity^19^. Our spatial transcriptomics analysis has demonstrated that spatial heterogeneity can result in distinct tumor immune microenvironments, leading to resistance and recurrence to immunotherapy in liver cancer^20^. To fully utilize the wealth of information contained in the spatial data in the TME, we coupled an agent-based model (ABM) with our whole-patient QSP platform to formulate a spatial QSP model (spQSP). The spQSP framework has been used to simulate the dynamics of T cells and tumor cells spatially and qualitatively compared to multiplex imaging data for NSCLC and breast cancer^21–23^. Extending this model to combination immunotherapies of liver cancer and their effect on its complex TME requires modeling additional cell types.

In this study, we constructed an spQSP model to computationally simulate clinical trial with neoadjuvant nivolumab (anti-PD-1 ICI) and cabozantinib (multitargeted tyrosine kinase inhibitor) therapy for patients with advanced HCC^3^. Accumulating evidence supports the importance of immunosuppressive macrophages on immunotherapeutic outcomes^24^. Similarly, angiogenesis is a well-established pro-tumor process in many cancer types, especially in HCC, and is thus targeted by many anti-VEGF/R therapies^4^. Therefore, in this study we developed a new spQSP model tailored to combination therapies in HCC that includes macrophages in the TME. Additionally, we developed a novel modeling strategy to incorporate angiogenic module to reflect the anti-angiogenic effect of cabozantinib. Together, using this new computational model a virtual clinical trial is conducted that simulates both patient outcomes and spatially resolved molecular states of tumors. We benchmark our computational model by comparing the simulated state of the TME to high-dimensional spatial proteomics and transcriptomics data from post-treatment tumor resections in the original clinical trial^3,20,24^. Whereas the biospecimens for the neoadjuvant clinical trial were only obtained at the time of surgery, the spQSP model fully simulated the spatial molecular states of the tumors over time. Therefore, once verified we can leverage this virtual clinical trial platform to develop an immunosuppressive score and investigate the molecular causes as candidate mechanistic pre-treatment biomarkers in future experimental and clinical studies.

## Methods

### Spatial QSP (spQSP) of HCC

In this study, we leverage the robust framework from our spQSP models to incorporate novel macrophage and angiogenesis modules that model combination therapy of cabozantinib and ICI in HCC (Fig. 1). The spQSP HCC model is based on our previous model^21,23^. Mathematical equations for cell modules in the model are included in the supplement. Below we only describe new modules in this study. The complete C++ code for the model is available as described in the Data Availability Statement to ensure reproducibility.

**Fig 1.**
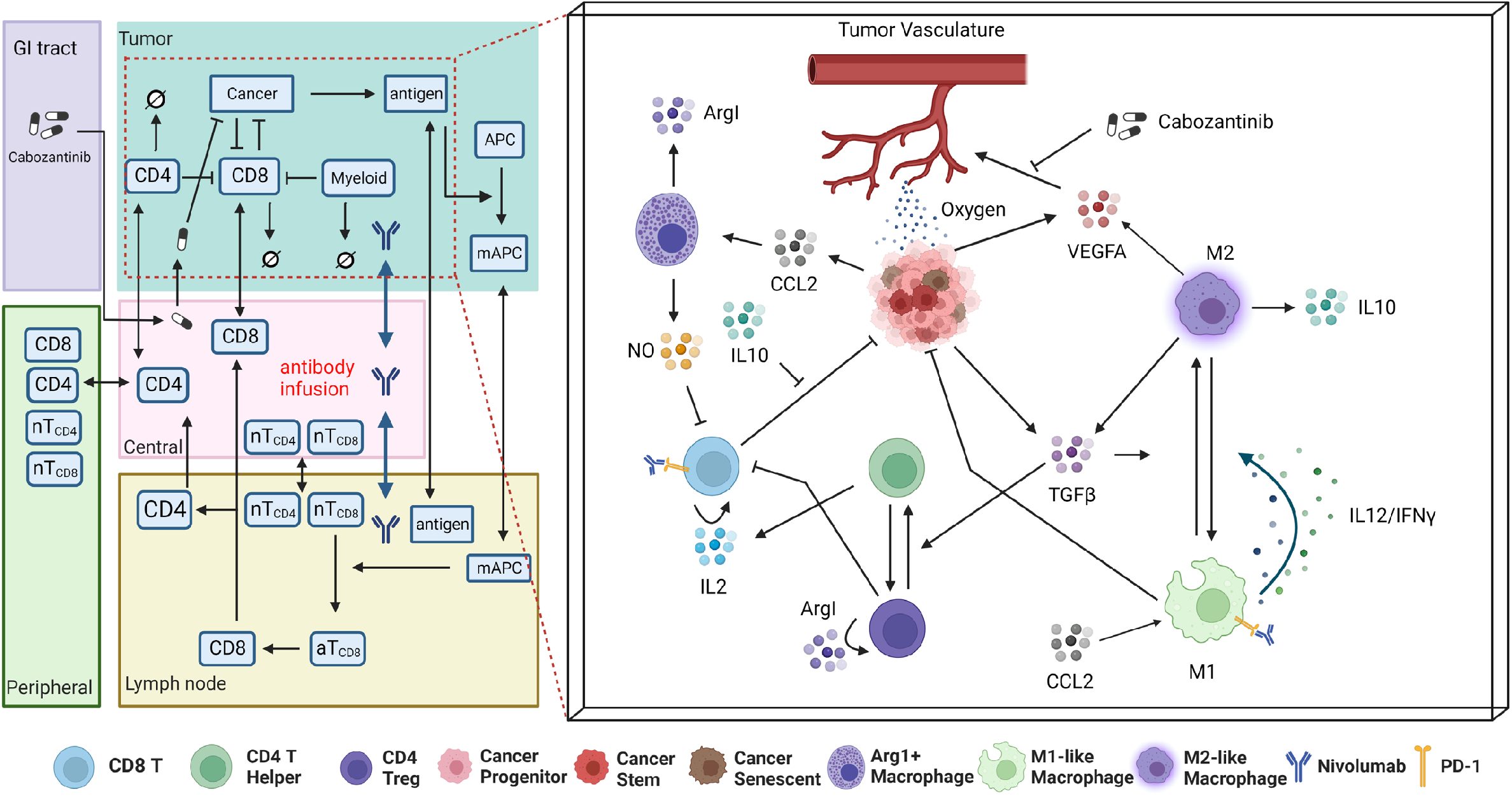
Schematic of the spQSP model for HCC immunotherapy integrating a systemic QSP model with a detailed ABM of the tumor and its microenvironment. Left: The QSP model simulates the systemic processes of T cell priming, immune cell trafficking, immune-cancer interactions, antigen collection and presentation, and pharmacokinetics and pharmacodynamics (PK/PD) of therapeutics. Right: Additional simulation of molecular components enabled by the ABM module of the tumor compartment (shown in red dashed box), which further models immune cell recruitment, cancer cell development and proliferation, immune cancer interactions, immune-checkpoint inhibition, and cytokine releasing and diffusion spatially.

### Agent-Based Model Setup

The Agent-Based Model (ABM) formulated in the study aims to reproduce spatial features extracted from both multiplexed image analysis and spatial transcriptomics sequencing. These datasets of the HCC tumors contain hundreds of millions of both cancer and immune cells, which is computationally unfeasible to simulate. To overcome this limitation, in the ABM we consider a flattened volume (6.5mm × 6.5*mm* × 200 *μm*), which is comparable with the size of histological specimens from HCC patients. Each voxel has dimensions 20 *μm* × 20 *μm* × 20 *μm*. Cells can move to their von Neumann neighborhood (6 voxels of adjacent neighbors) either randomly or guided by chemokine gradients; cells scan their Moore neighborhood (26 voxels of adjacent neighbors) for potential interactions.

The virtual patient cohort is generated by Latin Hypercube Sampling (LHS) based on estimated distributions^11^. Each set of model parameters is defined as a virtual patient, and each virtual patient cohort contains 15 patients in this study. For every virtual patient, an initial tumor diameter *D* is randomly generated, representing the pre-treatment tumor size. Fig. 2 presents the workflow of the spQSP model. The model is initialized with one cancer cell in the QSP module. When tumor diameter reaches *D*′ (*D*′ = 0.95*D*) in the QSP module, the ABM module is initialized. Both ABM and QSP modules are updated every Δt =6 hours. At a point τ, the ABM module is updated with QSP variables at t = τ. Next, both ABM and QSP modules are solved for t= τ+Δt. Then, ABM variables are updated back to the QSP, so that both modules are synchronized at t = τ+Δt. Treatments are applied when tumor diameter reaches *D*. Simulated spatial results at the end of the treatment are then compared with both multiplexed imaging and spatial transcriptomics analysis.

**Fig 2.**
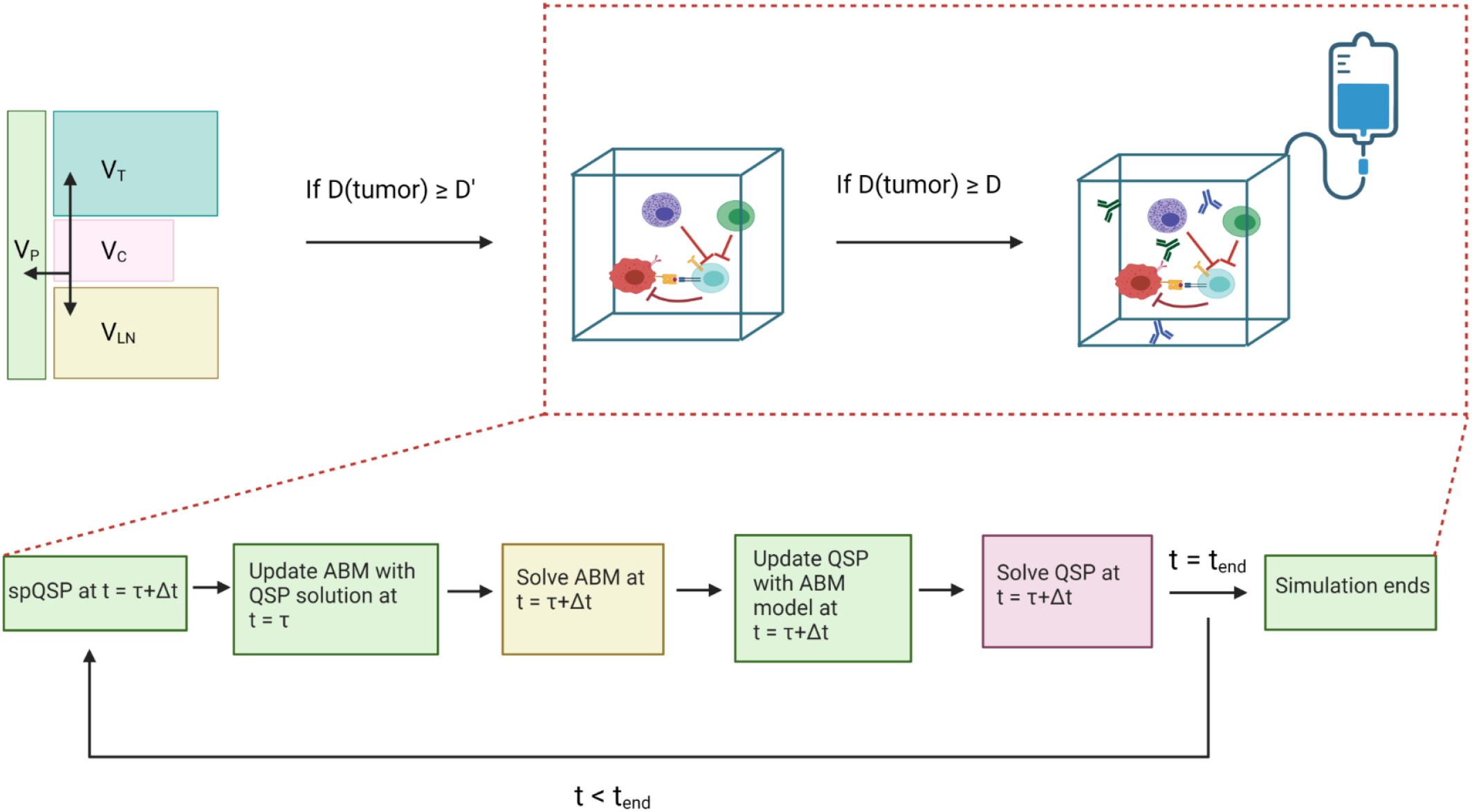
Top: The workflow of spQSP model. The ABM module is initiated when the tumor diameter reaches *D*′. Treatments are administered when the tumor diameter reaches *D* (*D*^′^ = 0.95*D*). Bottom: Synchronization between the QSP and ABM sub-model at each timestep during the simulation.

### Pharmacokinetics of Cabozantinib

In the phase 1b clinical trial (NCT03299946) on which the simulated patients from our spQSP model have molecular data for validation, cabozantinib is administered orally, 40 mg daily for a period of 8 weeks^3^. These values guide the timing of the simulated treatments in our model. Population based pharmacokinetic (PK) model for cabozantinib is based on clinical pharmacological data^25,26^. Previous work reported that the concentration-time profile of cabozantinib exhibits multiple peaks due to multiple absorption sites or enterohepatic recirculation or both. We assume that the pharmacokinetic model has multiple absorption sites along the gastrointestinal tract and is modeled as dual lagged (fast and slow) via first-order absorption and elimination processes. Following this cabozantinib is absorbed in the central compartment via first order absorption and diffuses to the peripheral, lymph node and tumor compartment. We assume nonlinear clearance of the drug from the central compartment. PK parameters are either taken from literature or optimized using the data reported in Nguyen et al. for healthy individuals^27^. PK parameters for cabozantinib are comparable for cancer patients and healthy volunteers^26^. Parameter optimization was performed using nonlinear least squares with trust-region-reflective method in Matlab (MathWorks, Natick, MA). The concentration of cabozantinib in the blood is characterized as:

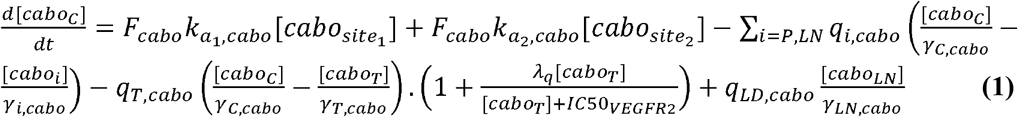

Here the first two terms on the right-hand side of the equation represent absorption of cabozantinib from the absorption sites in the GI tract to the central compartment, the third and fourth terms are the diffusive transport of the drug from the blood to the lymph node, peripheral and tumor compartment, the fifth term is the convective transport from the lymph node to the blood, and the last term is the clearance of cabozantinib from the central compartment. Cabozantinib interaction with VEGFR2 results in vascular normalization which increases transport rate of drugs from the blood to the tumor^28^; this has been incorporated by modification of the transport term for cabozantinib as well as for any drug in combination as depicted in the equation above. Cabozantinib concentration in the central (blood) compartment is shown in Extended Data Fig. 1.

### Pharmacokinetics of Nivolumab

The pharmacokinetic model is modified from our previously published QSP model on HCC^18^. Nivolumab (240mg) is injected intravenously into the central (blood) compartment every 2 weeks. The concentration of nivolumab in the central compartment is modeled as:

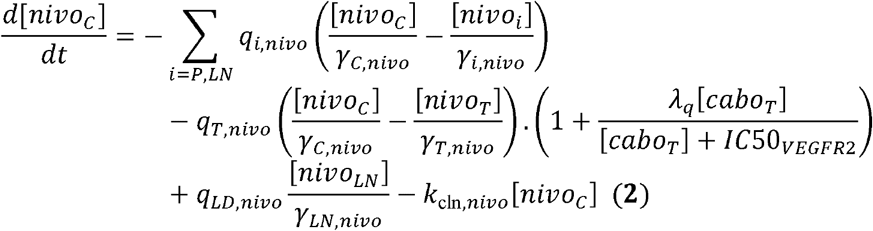

Terms for diffusive transport from central compartment to peripheral, tumor, and lymph node compartment are similar to Eq. 1, replaced with nivolumab-specific parameters. The parameters were initially calibrated under non-small cell lung cancer settings^29^. Sové et al. further optimized pharmacokinetic model in the context of HCC^18^.

### Spatial proteomics and transcriptomics analysis of neoadjuvant HCC

HCC samples were surgically resected as part of the clinical trial (NTC03299946) for neoadjuvant cabozantinib and nivolumab for patients with advanced stage HCC^3,20^. From 12 post-treatment FFPE surgical samples, we selected 37 tumor region cores to construct a tissue microarray (TMA). Spatial proteomics data were then obtained using the Hyperion Imaging System (Fluidigm, South San Francisco, CA)^24^. The same surgical specimen was also immediately embedded in optimal cutting temperature (OCT) compound and immediately frozen. A 10 μm cryosection was placed on a Visium Gene Expression slide (10x Genomics, Pleasanton, CA) for spatial transcriptomics analysis.

### Latent space identification using CoGAPS

Each spatial transcriptomics sample data are filtered to remove low quality spots and log2 normalized. The CoGAPS algorithm is applied on the preprocessed spatial transcriptomic sample (CoGAPS version 3.5.8)^30^ to obtain latent patterns associated with distinct cellular phenotypes. The output of CoGAPS factorization has two parts: an amplitude matrix and a pattern matrix. The amplitude matrix contains gene weights, and the pattern matrix contains spots weights associated to each pre-defined latent feature (i.e., pattern, total features = 15). The cell type of each pattern is identified by high weight genes in the amplitude matrix^31^.

### SpaceMarkers analysis to identify markers of cell-cell interactions

The SpaceMarkers algorithm is designed to identify molecular changes occurring due to the interactions between two distinct cellular phenotypes. The algorithm inputs an expression matrix of the spatial transcriptomic sample, the composition of cellular phenotypes inferred from the pattern matrix from CoGAPS, and a pair of patterns (*p*_1_, *p*_2_) in which to evaluate interactions as inputs^32^. The algorithm identifies spatial regions called hotspots that contain cells associated with both *p*_1_ and *p*_2_, defined as interacting regions. Using the differential expression model of SpaceMarkers, a Kruskal-Wallis test is then used to compare gene expression within the interacting regions relative to other regions. In spQSP outputs, we replace expression matrix with the simulated cytokine concentration of each voxel. Because the cell types are known *a priori* in the computational model, we also replaced the pattern matrix with a *n* × *m* cell matrix, where *n* is the number of cells and *m* is the number of cell types. SpaceMarkers identifies cellular hotspots for each cell type using outputs from spQSP model, and changes in cytokine expression using the SpaceMarkers differential expression mode.

## Results

### Virtual clinical trial of immunotherapy mirrored clinical correlatives in phase 1b neoadjuvant clinical trial

This study develops a spQSP model to conduct an in silico virtual clinical trial to investigate the spatial landscape of tumor microenvironment in HCC during cabozantinib and nivolumab combination therapy. Fig. 1 illustrates the extensions from our previous modeling framework to study the spatial distribution of cancer cells and immune cells in triple-negative breast cancer (TNBC) and non-small cell lung cancer (NSCLC)^21–23^ to model the more complex microenvironment of HCC in the present study. Specifically, we added spatially resolved computational modules to simulate macrophages, vasculature, and oxygen delivery. Clinical outcomes can be assessed from the model simulations by following tumor cell content. A pathological response is defined as a 90% reduction in cancer cell counts.

Once pathological responses from the model were simulated virtually, we then compared the results to those observed in the phase 1b trial for patients with advanced stage hepatocellular carcinoma with the neoadjuvant administration of cabozantinib and nivolumab, with 15 patients enrolled (12 patients evaluable)^3^. To minimize the randomness in generating virtual patient cohort with small sample size and the stochastic effects of the ABM module, we generated four cohorts, each consisting of 15 virtual patients. The dosing strategy in our simulations is identical to the clinical trial (Fig. 3A). Out of 59 virtual patients, 19 (32.2%) achieved pathological response, with 95% confidence interval of 26.2% to 38.2% (Table 1, Extended Data Fig. 2A, B). This simulated response rate is consistent with the response rate observed in the phase 1b clinical trial.

**Fig 3.**
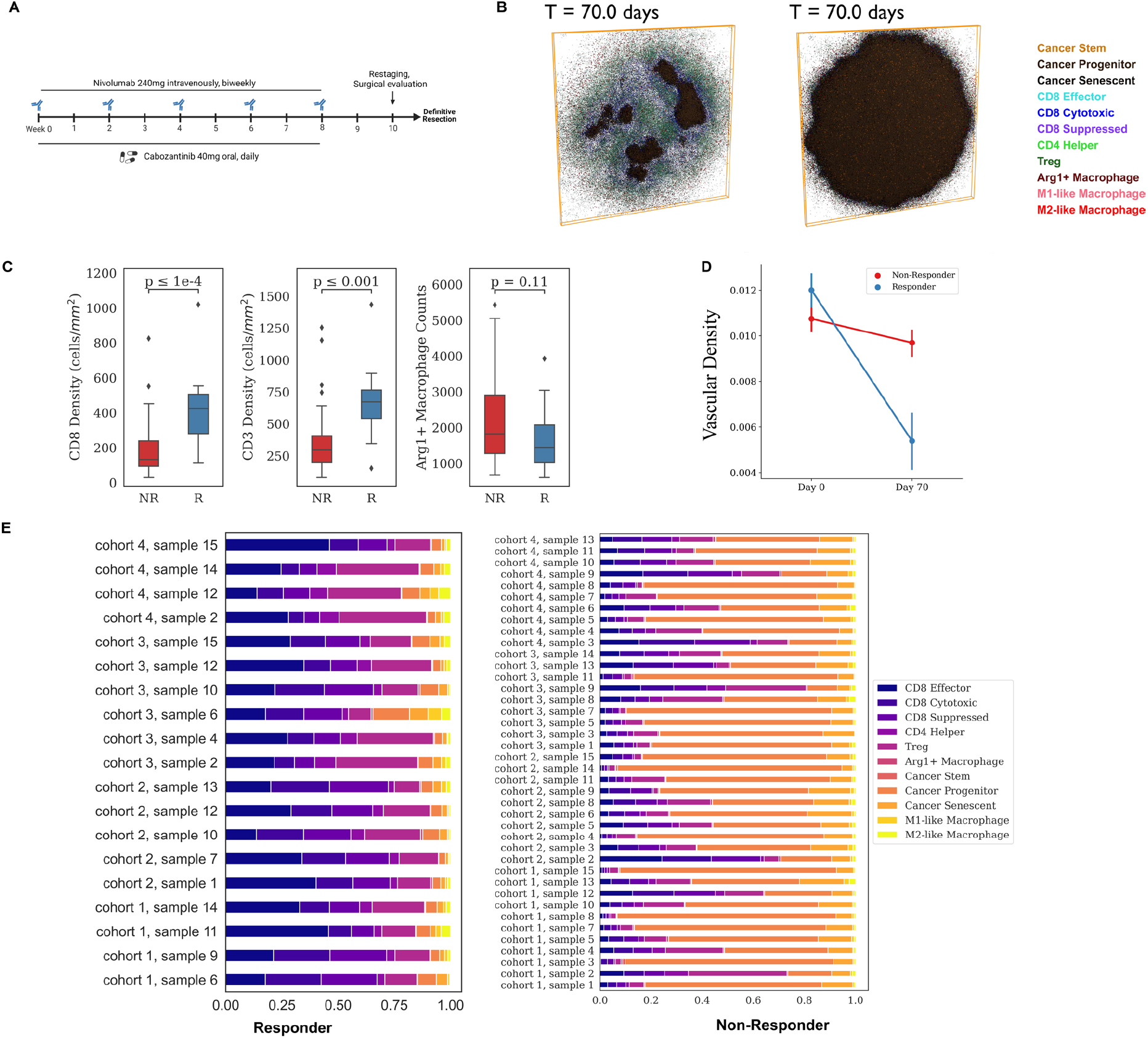
Results for the virtual clinical trial. A) Dosing strategy of nivolumab and cabozantinib in both the phase 1b HCC neoadjuvant clinical trial and spQSP virtual clinical trial simulations. Nivolumab (240mg) is injected intravenously every 2 weeks for 8 weeks. Cabozantinib is administered orally every day for 8 weeks. B) Two-dimensional cross section of the spatial distribution of cells in the tumor compartment from a representative simulation at day 70 for both responders and non-responders. Simulation movies for three-dimensional cellular states over time are provided in Supplement Movies. C) Quantitative comparison of CD8+, CD3+, and Arg1+ Macrophage in the stratified patient groups (responder: n=19 vs. non-responder: n=40) at day 70. D) Longitudinal dynamics of average vascular density in the ABM sample of two groups of patients (R vs. NR). E) Cell composition in the ABM model outputs at day 70, grouped by treatment outcomes.

The spatial resolution of the spQSP model enables us to simulate spatially resolved molecular data from the virtual clinical trial. By basing this simulation on the clinical trial, we have a unique opportunity to compare simulated molecular profiling data with real multi-omics datasets obtained from trial biospecimens. Spatially resolved virtual patient samples for a responder (R) and a non-responder (NR) can be obtained for all the molecular and cellular variables in the model and are shown in Fig. 2B and Extended Data Movie 1 and 2. The model outputs involve two parts: cellular output and molecular output. The cellular output includes the coordinates of each cell, along with its predefined cell type and state in the 3D space. The molecular output carries cytokine concentration of every voxel. List of cell types and cytokines in the model is presented in Fig. 1. The model is capable of fully resolving each of these measures in three-dimensional space. Slices across the simulation are used to summarize two-dimensional measures that can be compared to the molecular profiling data obtained in the clinical trial.

Based on our 2D simulated results, we observed a significantly higher density of CD8+ T cells in responders compared to non-responders (R: 407 ± 199 *cells*/*mm*^2^ vs. NR: 180 ± 155 *cells*/*mm*^2^). These values are comparable to clinical data (R: 493 ± 312 *cells*/*mm*^2^ vs. NR: 182 ± 177 *cells*/*mm*^2^). Similarly, we found a similar density of CD3+ cells between the simulated results (R: 657 ± 263 *cells/mm*^2^ vs. NR: 363 ± 261 *cells/mm*^2^) and clinical data (R: 773 ± 400 *cells/mm*^2^ vs. NR: 298 ± 252 *cells/mm*^2^) (Fig. 3C). Additionally, we observed that the non-responder samples had higher counts of Arg1 secreting macrophages (corresponding to hazard macrophages in Mi et al.^24^), although statistically insignificant, compared to the responder samples (Fig. 3C). To validate our simulation, we compared the vascular volume fraction (*V*_*vas*_) with the relative density of CD34 positive cells measured by Chebib et al^33^. The simulation yielded a range of 0.01 to 0.013, while the experimental measurement was 0.015^33^. Furthermore, when comparing the pre-treatment and post-treatment results in our simulation, we observed a decrease in *V*_vas_ for both responder and non-responder samples.

Ho et al. analyzed a paired pre-vs. post-treatment analysis using Nanostring PanCancer Immune Profiling panels, a multiplexed bulk transcriptional profiling technology^3^. Post-treatment multiplexed transcription data also revealed downregulation of endothelial marker CD31 and CDH5 after the treatment compared to pre-treatment results^3^. Simulation results indicate responders are observed with lower vascular *V*_*vas*_ compared to non-responders (Fig. 3D), which is in agreement with the results from another clinical trial for patients with advanced stage HCC treated with atezolizumab and bevacizumab^34^. Fraction of immune cells, including T cell and Arg1 negative macrophages (refer as macrophage), is higher in responder samples than the non-responder samples on Day 70 (Fig. 3E).

### Spatial metrics of cellular phenotypes define an immunosuppressive score that predicts clinical responses

One of the most important goals of constructing the spQSP model is to recapitulate not only bulk measures or population means in cells, but also the spatial characteristics from the unique spatial proteomic and transcriptomic profiling *in situ* in the surgical biospecimens. Our recent digital pathology study analyzing the spatial proteomics data from this study found the proximity between CD8+ T cell and arginase 1 positive (Arg1+), CD163 negative macrophage (defined as hazard macrophage) as a notable feature in non-responder samples^24^. For every CD8+ T cell, we denote *d*_1_as the center-to-center distance to its closest CD4+ T cell, and *d*_2_ as the center-to-center distance to its closest Arg1+ macrophage. Our spatial metric, immunosuppressive Score, is defined as 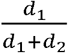 (Fig. 4A). To mimic the imaging mass cytometry (IMC) data from Ho et al. quantified with the immunosuppressive Score by Mi et al.^3,24^, we sectioned the 3D simulation result at y=5 position (i.e., in the middle of the 200*μm* slice) and generated 2D simulated imaging mass cytometry (IMC) data on both Day 0 and Day 70. The cancer cell region shrank by at least 90% on Day 70 while the tumor landscape remained unchanged in the non-responder sample (Fig. 4B, Extended Data Movie 3 and 4). The immunosuppressive Score is significantly reduced in responder samples compared to non-responder samples (Wilcoxon rank sum test *p* = 8.1 × 10^-4^), which is in agreement with the IMC studies (Fig. 4C). However, at pre-treatment stage, we observed smaller difference in immunosuppressive Score between responders and non-responders (Wilcoxon rank sum test *p* = 0.3).

**Fig 4.**
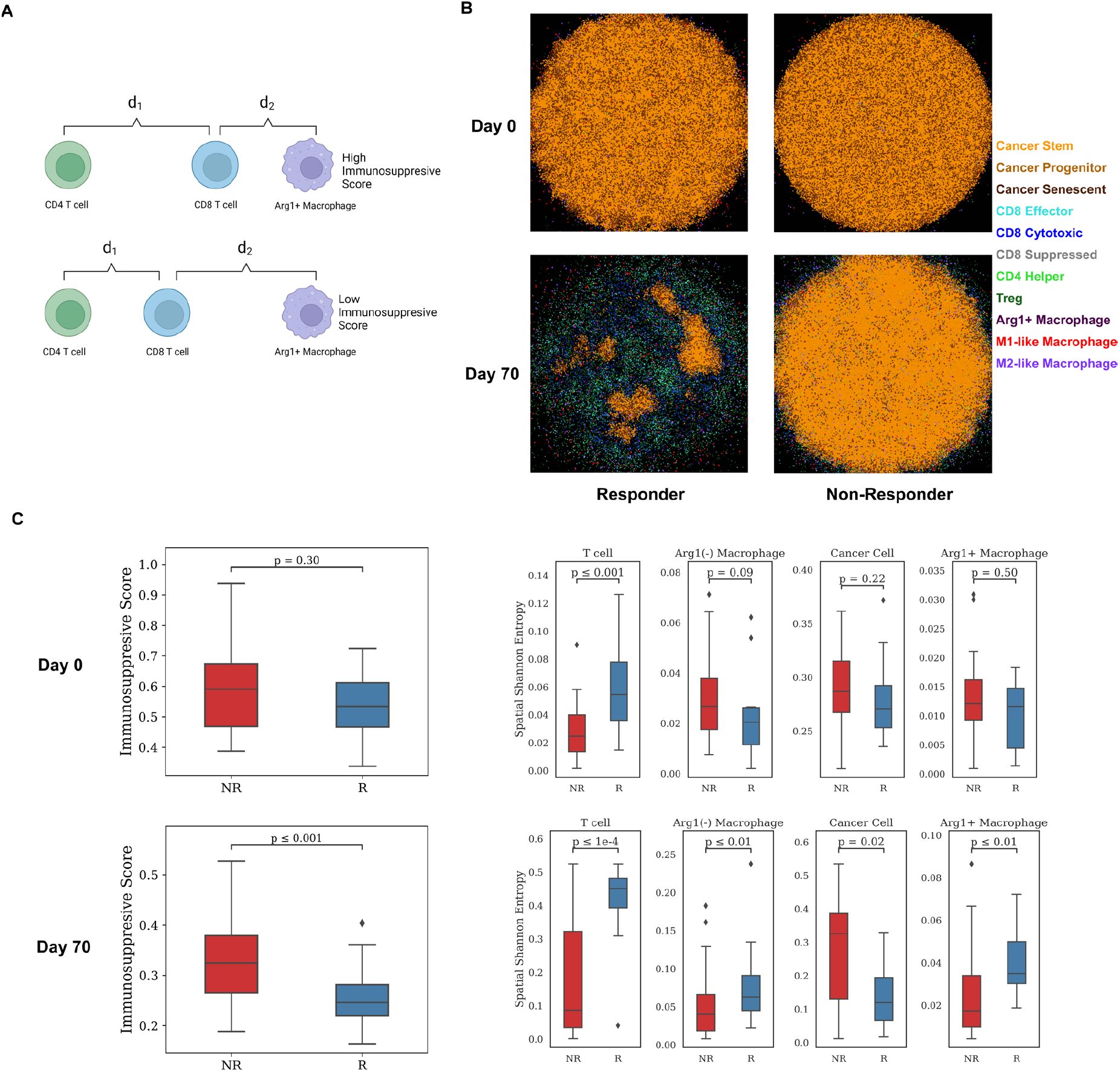
Spatial metrics summarized from model outputs. A) Schematic illustrates the definition of an Immunosuppressive Score. For each CD8+ T cell, *d*_1_ is defined as the distance to its closest CD4+ T cell, and *d*_2_ is denoted as the distance to its closest Arg1+ Macrophage. The Immunosuppressive Score is defined as 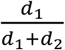. B) Simulated multiplexed imaging data used for calculating spatial metrics for responder and non-responder, respectively. Each sample is taken at y = 100μm. C) Spatial metric calculations based on simulated multiplexed imaging data of 60 virtual patients’ simulation. Left: Immunosuppressive Score calculated on per-cell basis, grouped by treatment outcome. Right: Spatial Shannon’s Entropy calculated for T cell, Macrophage, Cancer cell, and Arg1+ Macrophage in the simulated data at Day 0 and Day 70.

Our previous studies applied Shannon’s Spatial Entropy (SE) to multiplexed imaging analysis for HCC to quantify diversity and dispersion of various cell types in the TME^24^. The HCC study uncovered elevated SE for T cell, macrophages, and specifically Arg1+ macrophages in responder samples. Similar results were obtained in our simulations. SE for T cells, macrophages, and Arg1+ macrophages are higher in responder samples, while SE for cancer cells is increased in non-responder samples (Fig. 4C). At the beginning of the simulation, we observed higher T cell SE in responders, which provides a potential spatial biomarker for future studies. The analysis shows wider dispersion of immune cells in tumors of responders but more extensive cancer cell distribution in the non-responders in both simulation and clinical results.

### Simulated cytokines match patient spatial transcriptomics data suggesting tumor vasculature and TGFβ overexpression impact cancer and immune interactions

Our previous spQSP models and simulations have been qualitatively validated by multiplexed spatial proteomics data. These assays used pre-specified panels of proteins, often designed to resolve the cellular composition of tumor samples that can be compared to the simulated virtual tumors. The availability of whole transcriptome spatial data for the HCC clinical trial allows verification of spatial distribution of cytokines and cell types that are not profiled by multiplexed proteomics data. In addition, our new algorithm SpaceMarkers can further model molecular changes from cell-to-cell interactions^32^, providing an additional opportunity to validate the molecular regulatory programs in the computational model.

To verify the molecular layer of the spQSP platform, we identify regions of cellular co-localization using the SpaceMarkers algorithm in the same 2D region that we analyzed in the previous section. In the simulated results, the cancer region, CD8+ T cell region, and their interacting region are spotted in the responder sample (Fig. 5A, Extended Data Fig. 3). To our knowledge, this is the first spatial tumor model compared with both spatial transcriptomic data at molecular scale and multiplexed imaging data at cellular scale. To evaluate the stochasticity of the spQSP model, we repeated the simulation of one virtual patient five times. Stochasticity has little impact on the treatment outcomes (Extended Data Fig. 4). However, interaction regions were only identified for 3 replications using SpaceMarkers (Extended Data Fig. 5). Elongated cancer regions were observed for replicates 4 and 5. Therefore, future investigations must evaluate the impact of tumor shapes on identifying hotspot regions.

Within the simulation with an interaction region between cancer and immune cells, vascular density and TGFβ are overexpressed between CD8+ T cells and cancer cells. VEGFA is overexpressed in the cancer region, and IL2 expression is greater in the immune region (Fig. 5B, Extended Data Fig. 3). No immune region was identified in either simulated result or spatial transcriptomic data for non-responders due to cancer cell dominance in the TME, limiting our ability to infer comparable molecular changes in these non-responders (Extended Data Fig. 6, 7). Pro-inflammatory cytokines, including IL2 and IFNγ, have higher expression in the simulated responder than in non-responder sample (Fig. 5C).

**Fig 5.**
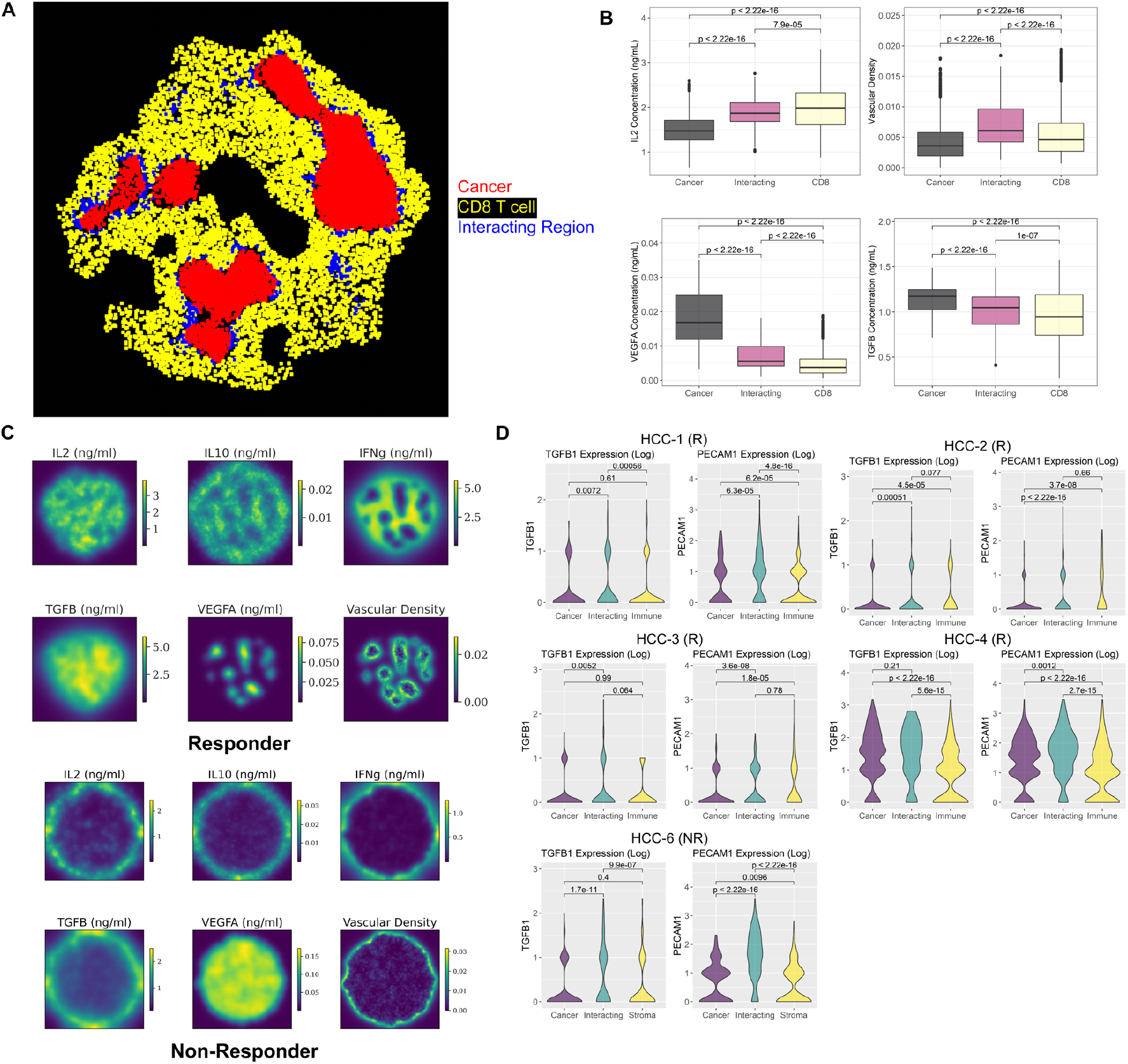
Spatial region identification and comparison with spatial transcriptomic analysis. A) The SpaceMarkers algorithm identified cellular hotspot regions of tumor and immune interactions in a simulated responder sample at day 70. B) Comparison of simulated cytokine concentration in the cancer cell region, CD8+ T cell region, and interacting region Identified in panel A using Kruskal-Wallis test. C) Simulated spatially resolved cytokine concentration and vascular density distribution for a responder and a non-responder sample at day 70. D) Expression of TGFβ and endothelial cell marker (PECAM1) in 5 spatial transcriptomic samples (4 responders and 1 non-responder) obtained from post-treatment surgical biospecimens in the phase 1b clinical trial. The DE model of the SpaceMarkers algorithm is applied to every sample to identify gene expression changes associated with interactions between cancer and immune cells.

We compared these simulated data to the SpaceMarkers interaction statistics for the real Visium spatial transcriptomics data obtained from the clinical trials biospecimens, with a focus on endothelial cell markers *PECAM1* (CD31) and immunosuppressive cytokine TGFβ. However, pro-inflammatory cytokines including IL2, IFNγ, and IL12 are not well captured in the spatial transcriptomic data (expressed in less than 3 spots per sample) and thus cannot be compared to the simulated data from our computational model. To connect these spatial patterns to patient response in the virtual clinical trial, we run SpaceMarkers on outputs of the virtual trial. Among 19 virtual responders in this cohort, interacting regions were identified in 14 virtual samples. In contrast, only 9 samples were observed with the interacting regions out of 40 virtual non-responders’ samples (Extended Data Table 1), which demonstrate that virtual patients with immune-cancer interacting regions tend to respond to the therapy (Chi-Squared Test: *p* = 3.1 × 10^-5^).

In real human spatial transcriptomic data, we also apply SpaceMarkers to identify regions of interactions between cancer and immune cells (Extended Data Fig. 8, 9). In simulation results, vascular density is significantly higher in the interacting regions (Fig. 5B, Extended Data Fig. 3). Analogously, *PECAM1* (CD31) is robustly overexpressed in the interacting region in all five spatial transcriptomic samples (Fig. 5D). Expression of other endothelial markers including *CDH5* and *CD34* further proved higher tumor vascular density in the interacting region (Extended Data Fig. 10). Concentration of TGFβ is increased in the interacting regions in some simulated samples while exhibiting no significant difference in other simulation results (Extended Data Fig. 3). In the spatial transcriptomics data, TGFβ is overexpressed in the interacting region in some samples (HCC-1, 3, 6) but other samples show no difference (HCC-2, 4) (Fig. 5D). Among 23 samples in simulated patients in the virtual clinical trial identified with an interacting region between cancer and immune cells, TGFβ overexpression is observed in 8 samples. On the other hand, 16 samples from simulated patients were found elevated vascular density in interacting regions between cancer and immune cells (Extended Data Table 1). Thus, spatial transcriptomics results for the spatial distribution of various cytokines and vascular density are in qualitative agreement with the data simulated by the spQSP model. Our simulation results suggest elevated tumor vasculature and TGFβ level in the interacting region of cancer and immune area, which is consistent with our spatial transcriptomic analysis.

### Proximity between CD8+ T cell and Arg1+ macrophage, cancer growth rate, and stem cell markers are identified as predictive biomarkers

After examining the spatial metrics at cellular and molecular resolution and comparing simulated post-treatment results with acquired TMA and spatial transcriptomics data, we analyze the simulated pre-treatment data to predict potential spatial and non-spatial biomarkers. Although these predicted biomarkers cannot be validated with current data because of the small sample size, they can provide insight for future clinical trial design (Fig. 6). As expected, high CD8+ and CD3+ T cell densities predict higher likelihood of responding to the therapy. Patients with fewer Arg1+ macrophage counts are also prone to respond to the therapy, which is in agreement with previous studies^35,36^. In addition, higher ratio between M1-like and M2-like macrophages (M1/M2) reflecting macrophage polarization status is associated with better response rate, since M2-like macrophages are one of the sources of TGFβ, an immunosuppressive cytokine. Spatial metrics show that higher distance between CD8 T cell and Arg1+ macrophage corresponds to higher response rate. The closer proximity between CD8+ T cell and Arg1+ macrophage makes CD8+ T cell more susceptible to exhaustion via paracrine signaling of both Arg1 and NO.

**Fig 6.**
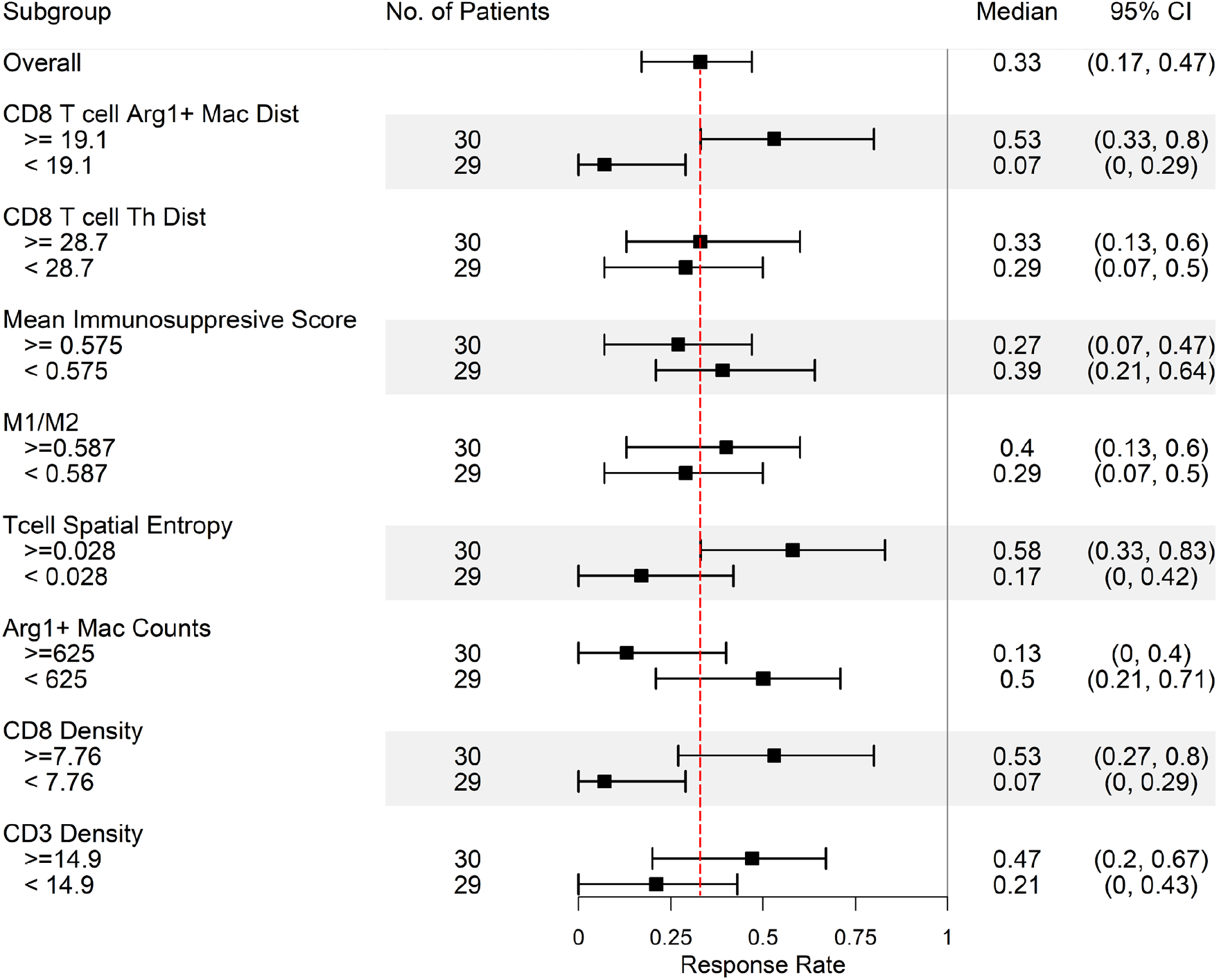
Biomarker identification at pretreatment stage. Application of the spQSP model for biomarker identification based on the pre-treatment composition of the HCC tumor microenvironment. Virtual patients are divided into upper half and lower half the day 0 values of 8 different features. Simulated median response rates (90% cancer cell reduction in the ABM model) at day 70 after treatment of every subgroup are computed along with 95% bootstrapped confidence intervals.

To investigate the impact of model parameters used to generate virtual patient cohorts, we performed the partial rank correlation coefficient (PRCC) sensitivity analysis in both QSP model and ABM. The cancer growth rate and initial tumor diameter are highly related to cancer cell counts by the end of the treatment (Fig. 7). The cancer growth rate is normally estimated from abundance of Ki-67 from the immunofluorescence data or expression of proliferation related marker in the transcriptomic data^37^. Both are strong predictors of therapeutic responses. In addition, the number of CD8+ T cell clones are associated with lower cancer cell counts, and studies have suggested that richer CD8+ TCR clones predict better response^38,39^. In contrast, even though a higher number of CD4+ clones give higher helper T cell counts, it also increases the infiltration of regulatory T cell which suppresses the cytotoxicity of CD8+ T cell resulting in less optimal treatment outcomes. Elevated helper T cell recruitment decreased the immunosuppressive Score. The recruitment rate of Arg1+ macrophage not only positively correlates with cancer cell counts at the end of the treatment, but also positively correlated with higher immunosuppressive Score.

**Fig 7.**
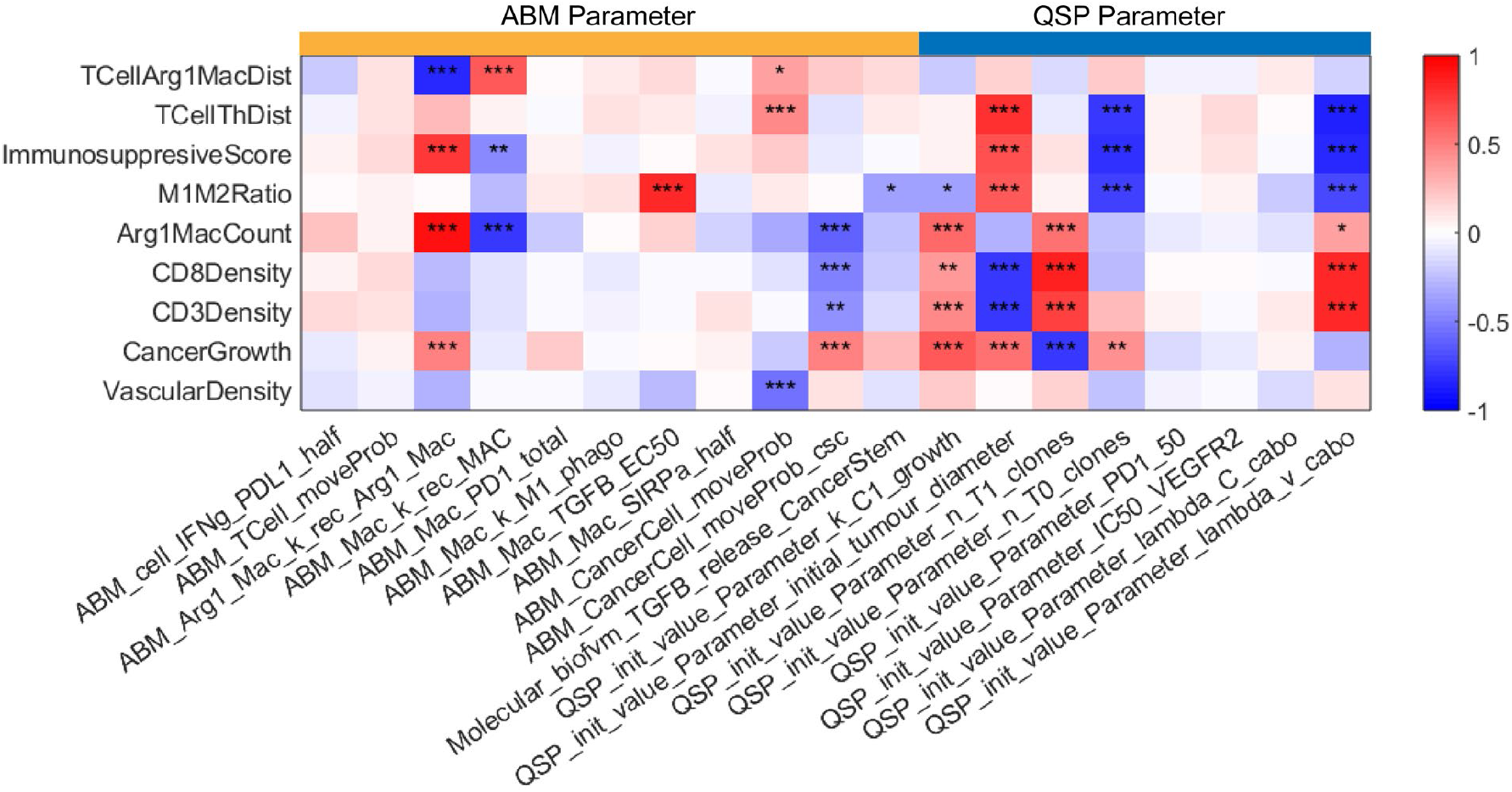
Sensitivity analysis. The sensitivity analysis of 19 model parameters (11 ABM parameters and 8 QSP parameters) using the PRCC method (*: p < 0.05; **: p <0.01; ***: p<0.001), measuring the partial rank correlation coefficient (ranging from -1 to 1) between the model parameters (each column) and output variables (each row). Detailed biological interpretation of all model parameters is included in the supplemental materials.

One observation from PRCC results is that high motility of cancer stem cells is associated with poor treatment outcomes while motility of progenitor cancer cells shows the opposite trend. Consistent with this observation, we note that our previous spatial transcriptomics analysis of the trial samples found enrichment of cancer stem cell markers within a region of low immune infiltration in the only patient with recurrence in the trial^20^. Cancer stem cells with higher migration rate form more aggressive tumor niche and prone to metastasis^40^. However, the metastasis compartment is not the focus of this study and requires additional extensions to our model in future work.

## Discussion

In this study, we developed a new virtual clinical trial framework by creating an spQSP model to analyze the clinical outcomes of a recent clinical trial in advanced HCC patients who underwent neoadjuvant therapy with nivolumab and cabozantinib. We utilized a compartmental QSP model to track tumor progression at the organ level and employed a coupled agent-based model describing cells and their interactions to monitor the dynamics of the tumor microenvironment with single-cell resolution. Previously, we demonstrated the integration of neoantigen profiles from single-cell RNA sequencing with our spQSP model to relate antigen homogeneity in tumor cells with therapeutic outcomes^22^. With the enrichment of spatial data, we now leverage spatial features from tumor biospecimen to evaluate the role of the TME on patient response in both virtual and phase 1b clinical trials. To enable this investigation in HCC immunotherapy, we developed a new spQSP model incorporating modules describing macrophage polarization and tumor angiogenesis to evaluate the impact of these processes on treatment outcome.

Spatial proteomics analysis at time of surgery simulated in the spQSP model and from IMC profiling of the phase 1b trial biospecimens enabled us to establish an immunosuppressive Score, indicating that the relative distance between T cells and Arg1+ macrophage in the tumor is linked to patient outcomes. Moreover, comparable analyses of spatial transcriptomics data revealed TGFβ overexpression in the interacting region between tumor and immune regions, which was consistent in both patient data and simulation outputs. While these assessments could be performed from the spatial molecular data in the trial samples directly, these biospecimens are only obtained from a single moment in time and may not fully reflect the dynamic changes within the tumor microenvironment over the course of treatment. To address this limitation, our computational model aims to simulate the dynamics of the tumor microenvironment throughout the treatment by calibrating it with the available spatial data. As a result, the computational model can simulate the spatial molecular state of tumors pre-treatment. We relate these simulated pre-treatment spatial data to propose CD8+ T cell and Arg1+ macrophage cell proximity as candidate spatial biomarkers of patient response. Additionally, we observed a significant association between stem cell motility and treatment outcomes in the virtual clinical trial. Although these candidate pre-treatment biomarkers require further validation in future clinical studies, they highlight the clinical value of our computational model to inform the design of clinical trial correlates and predict patient outcomes. Future work informing our model with patient-specific omics data will also enable personalized simulations, bridging the gap between clinical measurements, especially considering the limited opportunities for biopsy and resection in neoadjuvant trials.

The study is limited by the sample size of pathology samples we acquired from the clinical study. The spQSP model is built on 12 evaluable multiplexed imaging specimens (out of 15 patients) plus 7 out of 15 spatial transcriptomic data due to sequencing quality issues. The model might not be as robust as models built based on larger clinical trials. Nonetheless, we note that the high-dimensional spatial multi-omics profiling of this neoadjuvant trial provides an unprecedented wealth of data to test our spQSP model at both the cellular and molecular levels. In addition, our model is also limited by the number of cell types simulated. Future studies expanding the interactions with other cell types could provide a more comprehensive landscape in the TME using spQSP model. Since the spQSP model is highly modularized, additional cell modules generally do not require modifications of existing modules. Notably, our independent analysis of the spatial transcriptomics analysis of this trial show cancer-associated fibroblasts (CAFs) and extracellular matrix (ECM) components, such as collagen, fibronectin, and vimentin, predominantly in non-responder samples^20^. Studies found the immunosuppressive effect of ECM by physically blocking immune cells from contacting cancer cells, and ECM density is negatively correlated with T cell motility^20,41^. Clinical data reveal high density of B cells and tertiary lymphoid structures (TLS) correlated with superior prognosis^42,43^. The cause for forming TLS in some patients but not others is not yet clear, and the role of B cells in HCC seems to be underestimated. Antibody production and antigen presentation to T cells are two most well-known functions of B cell^44^. Incorporation of B cells and CAFs into the spQSP platform should help uncover suitable prognostic markers under various clinical settings.

To summarize, this paper presents an integrative model that combines multiscale continuous modeling and agent-based modeling approaches to capture the complexity of the HCC tumor microenvironment while balancing the number of model parameters. By integrating these models with neoadjuvant clinical trials, the simulations can be grounded in real-world patient outcomes and suggest novel pre-treatment biomarkers of patient response. Although a potential more complex computational models of the full high-dimensional cellular and molecular landscape of the TME of HCC accurately reflect human tumors, parameter fitting problems become more challenging, requiring more data for parameterization. To address this challenge, spatial metrics are used to define low-dimensional statistical similarities between simulated data and real clinical data, particularly in the context of stochastic agent-based models. For example, Hutchinson and Grimm presented an example of using pre-and post-treatment digital pathology data in combination with a simple two-dimensional agent-based model^45^. Other studies have employed neural networks to project image data onto lower-dimensional spaces, where the distance between real and simulated data in this space is used to measure similarity^46^. Since running ABM with partial differential equation (PDE) solvers is highly time consuming, machine learning based (ML-based) surrogate model are proposed^47^. The surrogate model learns the behavior of ABM model and predicts the model outcome given the parameter input to reduce computational cost. However, the outcomes from the ML-based surrogate are sets of abstracted spatial metrics rather than exact location of every agent limiting the ability to calibrate with real world data as in the mechanistic parameters of the spQSP model in this study. In all cases, data assimilation methods that formally embed patient datasets into these computational models may further enable extending these models from virtual cohorts to predictions of outcomes in individual patients^48,49^.

## Supporting information

All supplemental Figures

Supplemental equations

Model parameters

## Acknowledgements

The authors thank Andrew Ewald and Phuoc Tran for feedback.

## Data Availability Statement

The authors confirm that the data supporting the findings of this study are available within the article and the Supplement. C++ code for model generation and virtual clinical trials can be found at https://github.com/popellab/SPQSP_IO_XXXX. [The code will be made available to reviewers on GitHub. The code will be made public on GitHub and assigned a Digital Object Identifier by Zenodo upon acceptance].

