## Supplementary material for "Informing virtual clinical trials of hepatocellular carcinoma with spatial multi-omics analysis of a human neoadjuvant immunotherapy clinical trial": All supplemental Figures

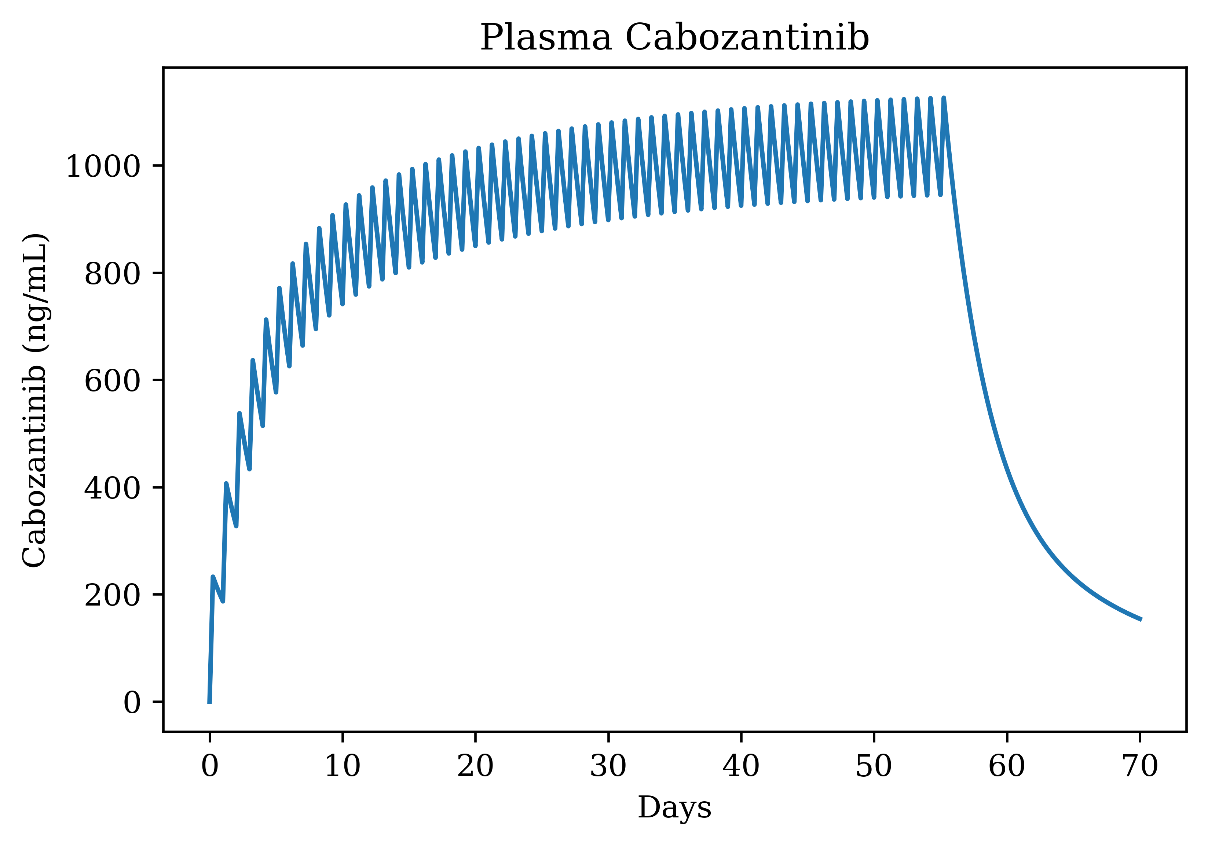

Extended Data Fig. 1. Plasma Cabozantinib concentration (ng/mL) during simulation (70 days).

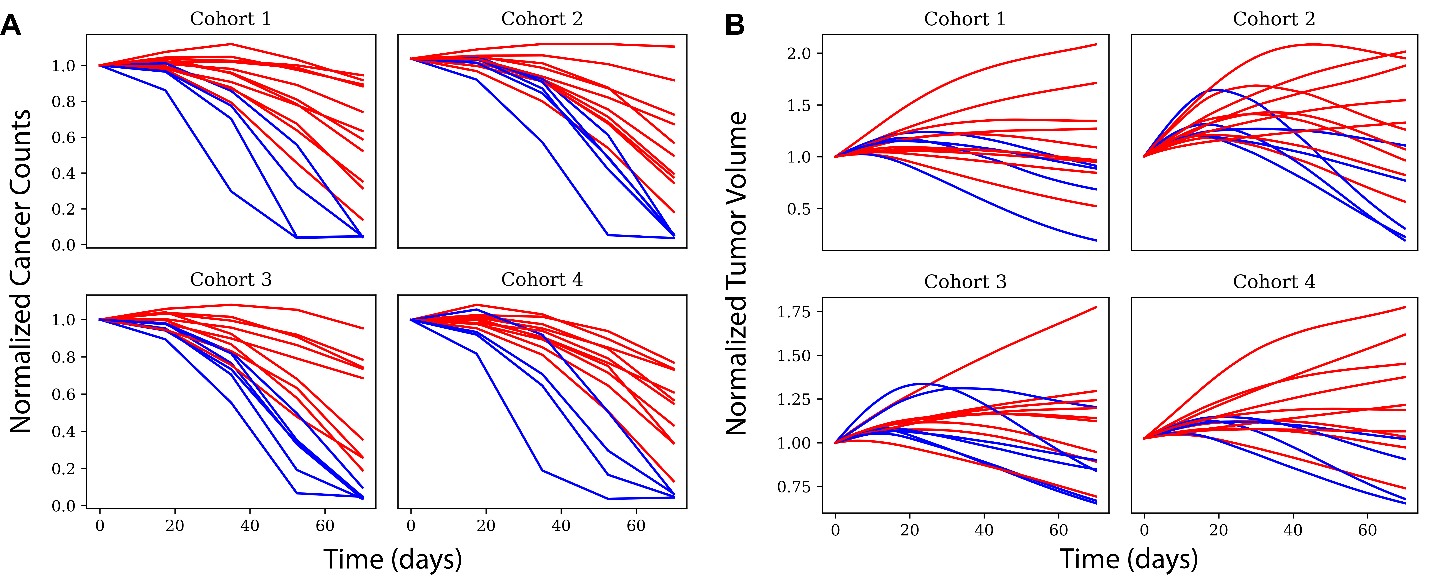

Extended Data Fig. 2. **Tumor trajectory over virtual treatment.** A) Longitudinal dynamics of normalized cancer cell counts in the ABM sub-model under combination therapy. B): Change of relative tumor volume in the QSP model over the course of treatment under combination therapy.

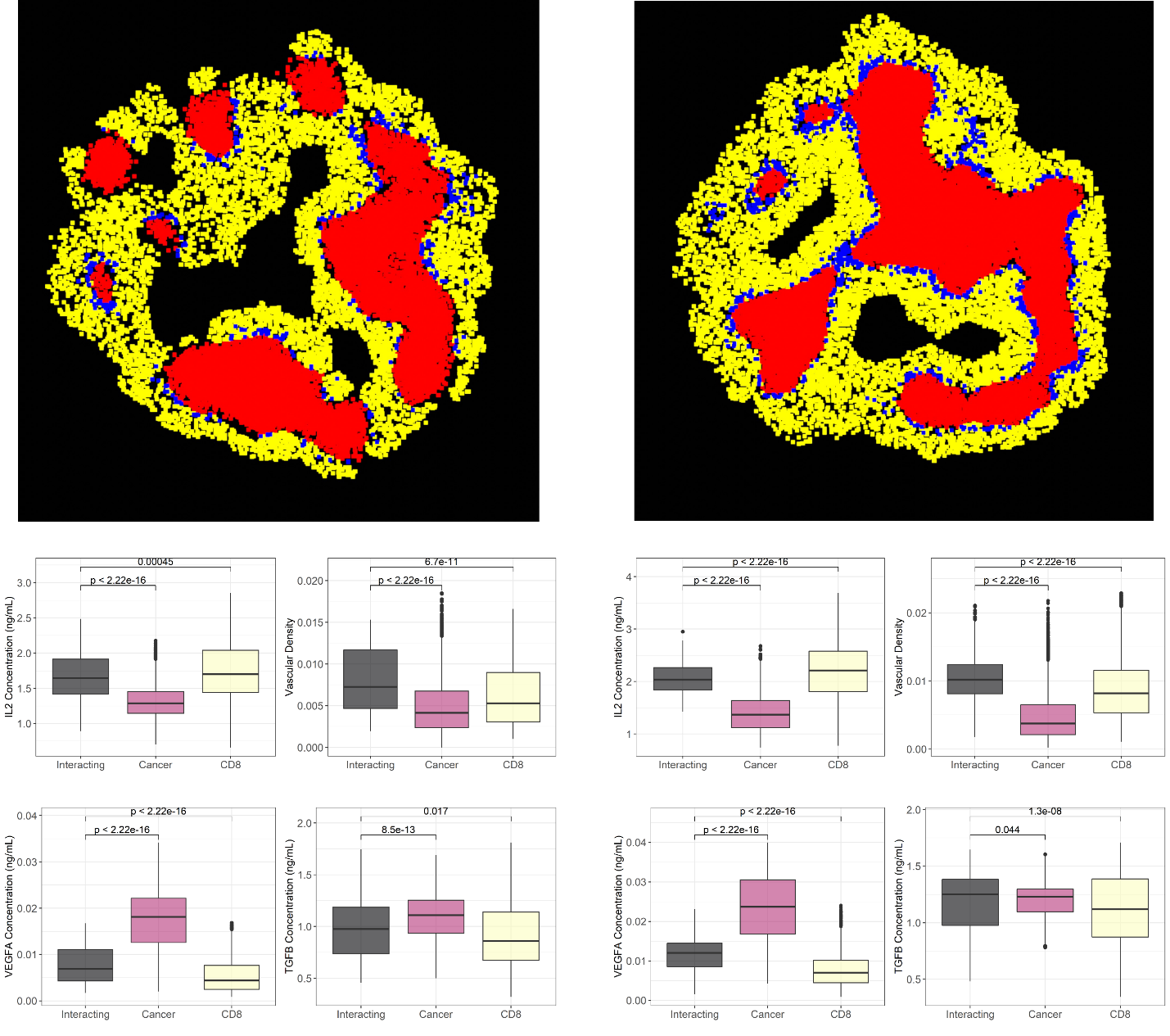

Extended Data Fig. 3. **Additional simulation samples for two virtual responders.** Top: Identified cellular hotspot regions. Red: Cancer; Yellow: CD8+ T cell; Blue: Interacting region. Bottom: Corresponding cytokine expression and vascular density profiles across three regions.

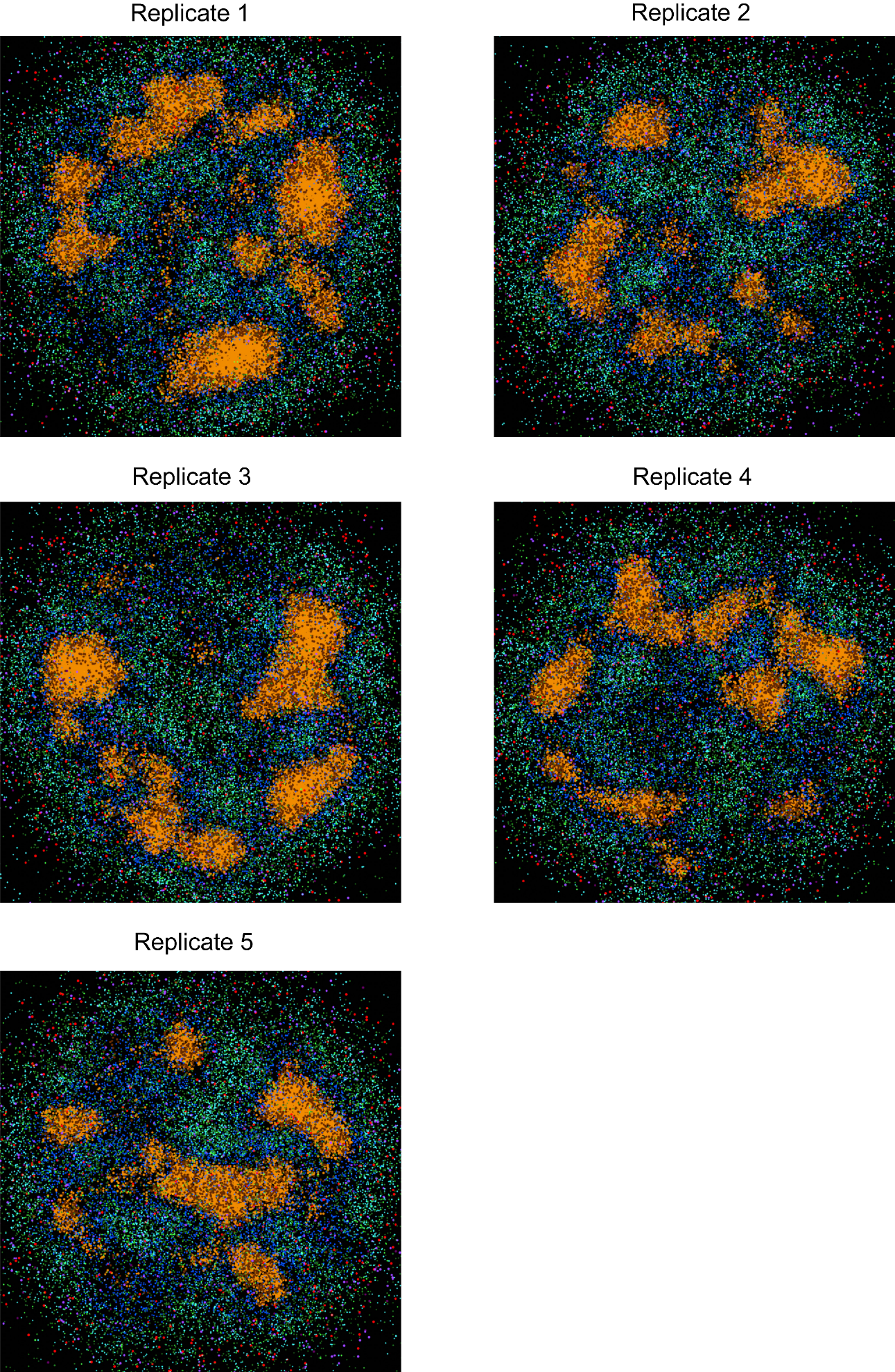

Extended Data Fig. 4. **Repeated simulation to check the impact of stochasticity in spQSP model.** Spatially resolved simulation outputs at Day 70 for the same virtual patient. Simulations are repeated 5 times. The color schematic is identical to Fig. 4b.

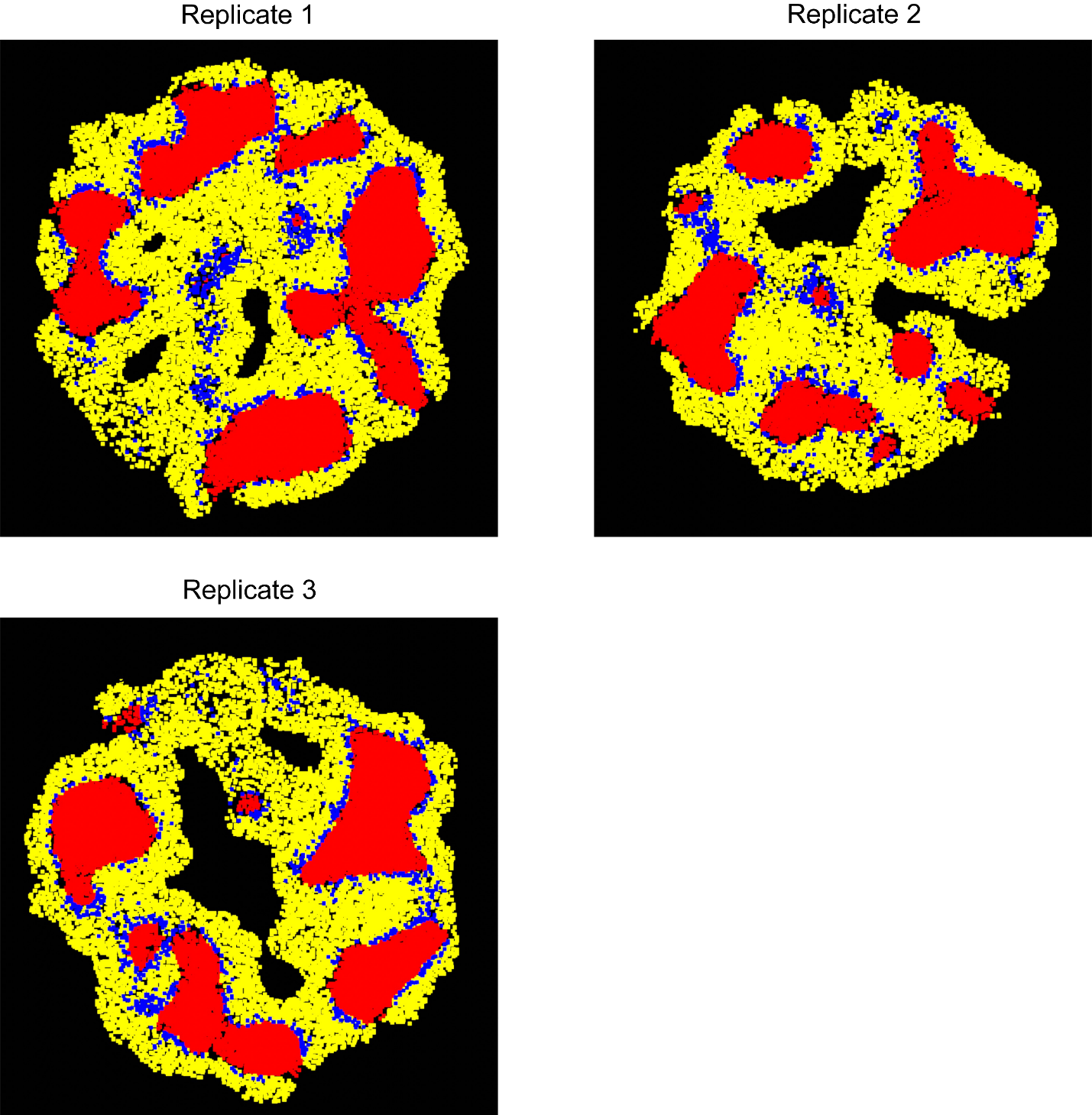

Extended Data Fig. 5, **Hotspot identification using SpaceMakers**. SpaceMarkers is applied on the 5 repeated simulations for the same virtual patient. Interactions regions for Replicates 4 and 5 were not identified. Red: Cancer; Yellow: CD8+ T cell; Blue: Interacting region.

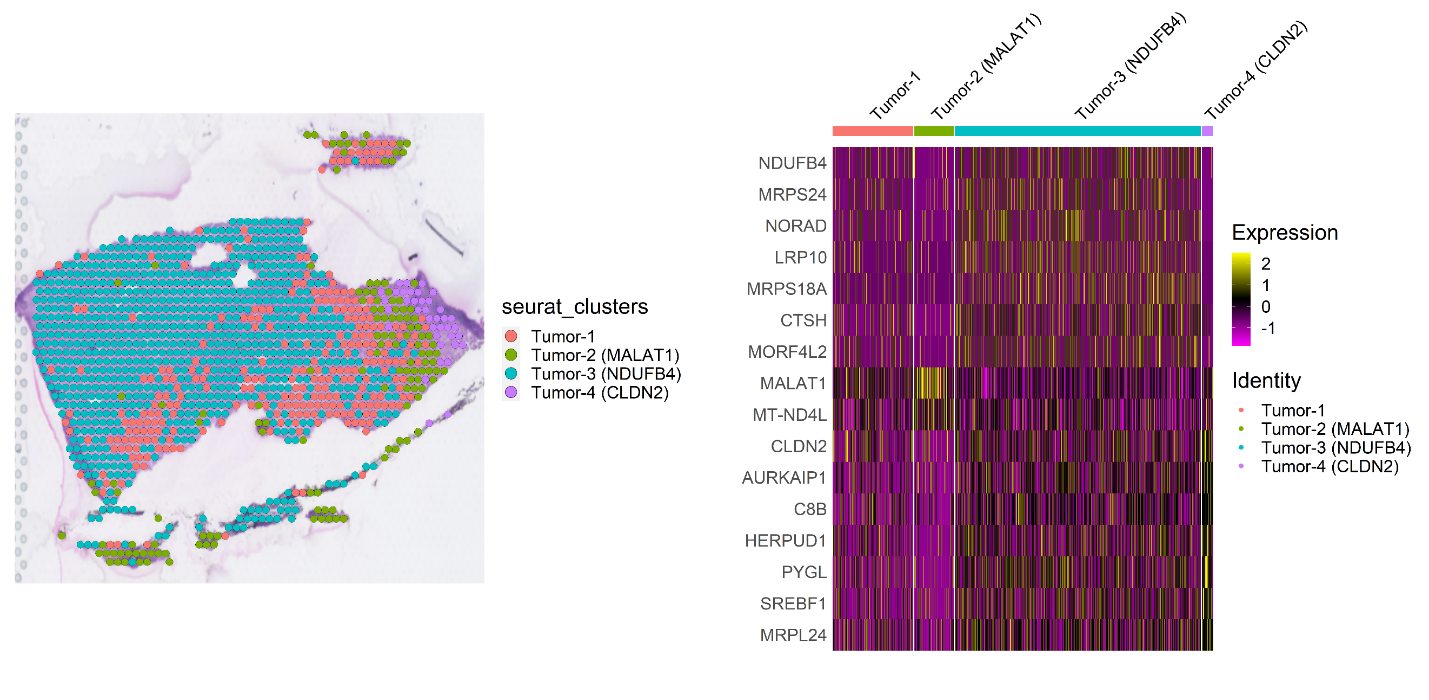
 Extended Data Fig. 6. **Region clustering for patient HCC6-NR.** Histological composition of the specimen using unsupervised clustering (Left panel). The clusters are identified based on the top variable genes (right panel). Results contains no immune region, which is not suitable for SpaceMarkers analysis.

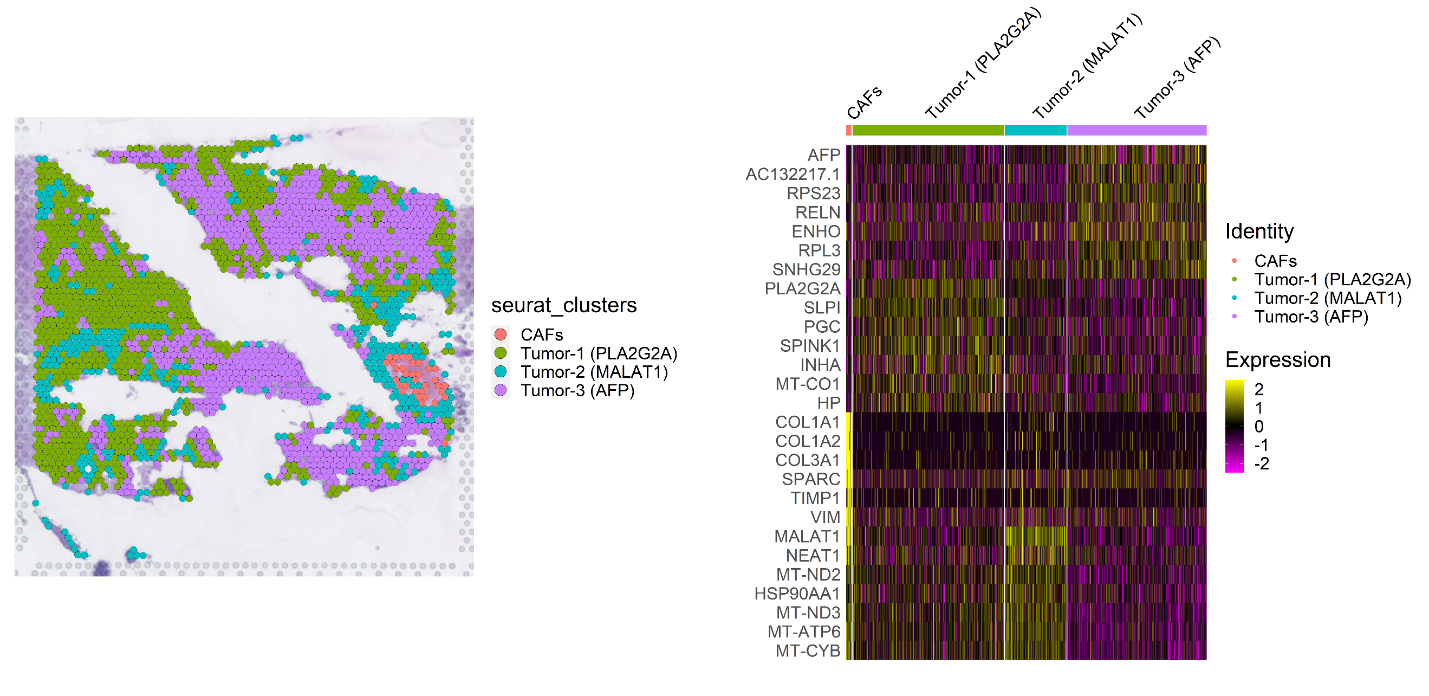
 Extended Data Fig. 7. **Region clustering for patient HCC7-NR.** Histological composition of the specimen using unsupervised clustering (Left panel). The clusters are identified based on the top variable genes (right panel). Results contains no immune region, which is not suitable for SpaceMarkers analysis.

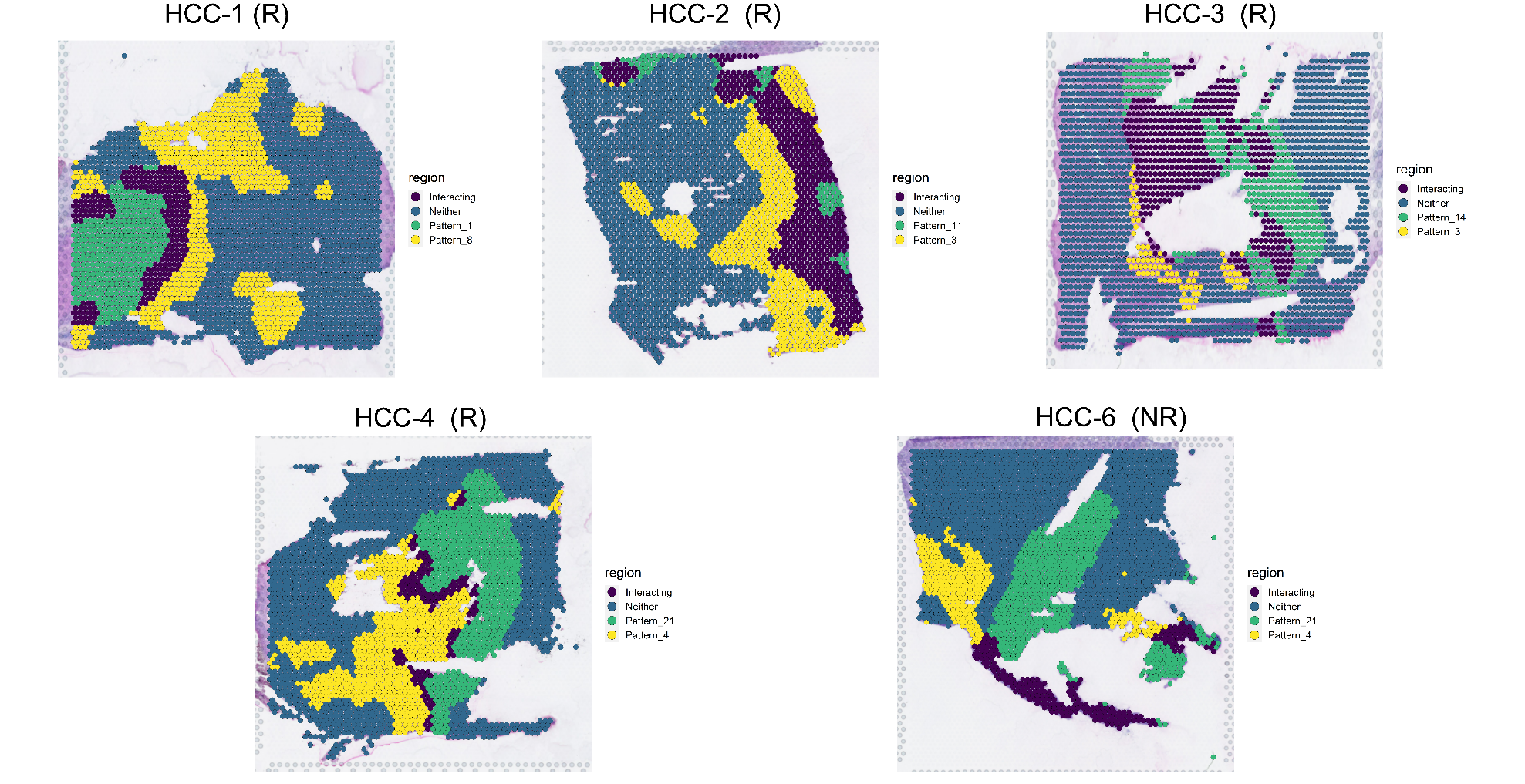

Extended Data Fig. 8. **Hotspot identification results using SpaceMarker.** CoGAP analysis is first applied to all spatial transcriptomic data samples. Then we used Pattern Spotter to identify hotspot regions of each pattern for every sample. Finally, we used expression profile of each pattern to infer associated cellular regions

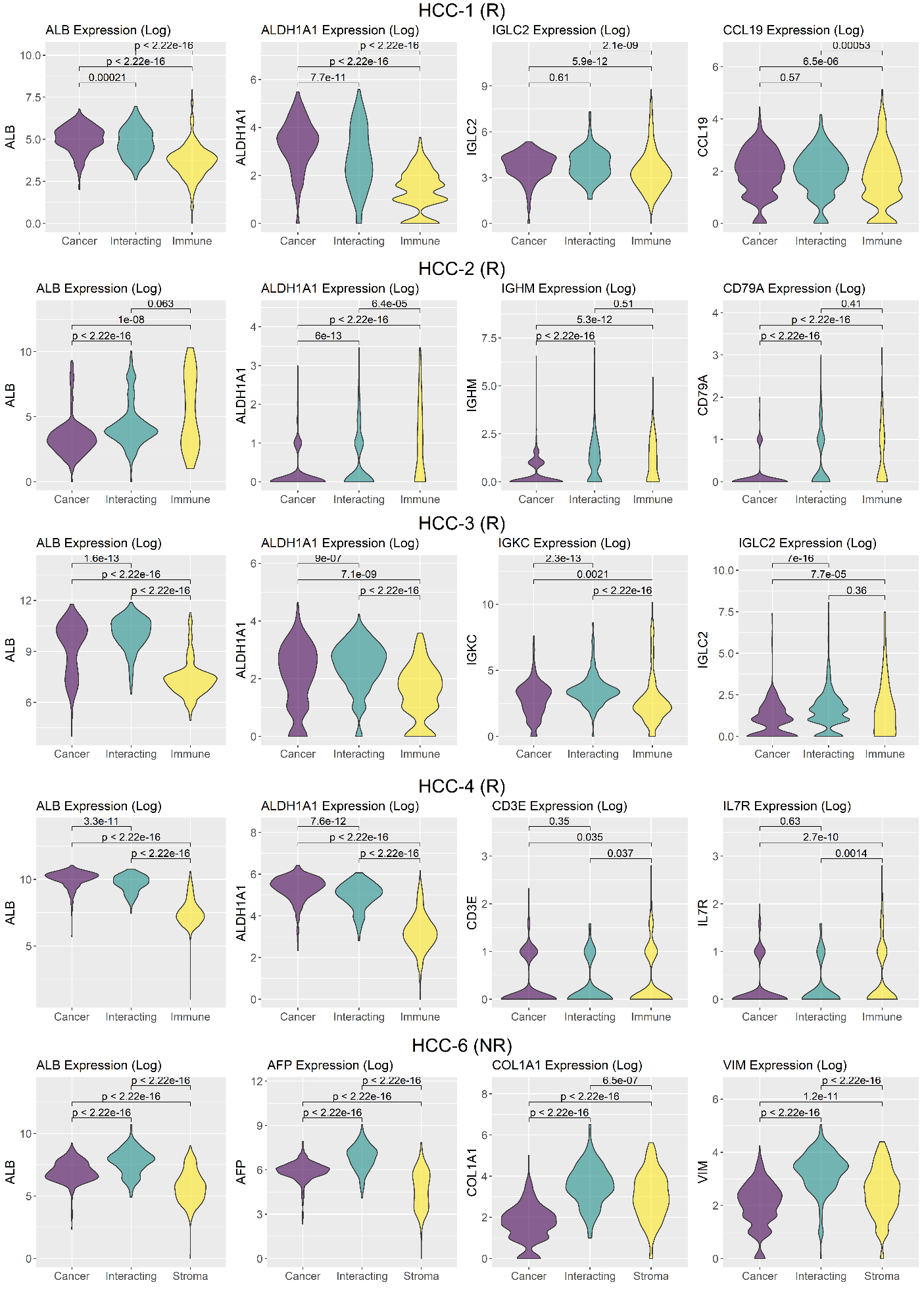

Extended Data Fig. 9. **Pattern identifications using by signature genes.** Biological phenotypes of each CoGAPS pattern are identified by the expression levels of tumor signature genes (*ALB, AFP, ALDH1A1*), immune signature genes (*IGLC1/2, CCL19, CD3E, IL7R, IGHM, and CD79A*), and stroma signature genes (*COL1A1, VIM*). The violin plots illustrate expression levels of signature genes in cancer regions (purple), interacting regions (cyan), and immune regions (yellow).

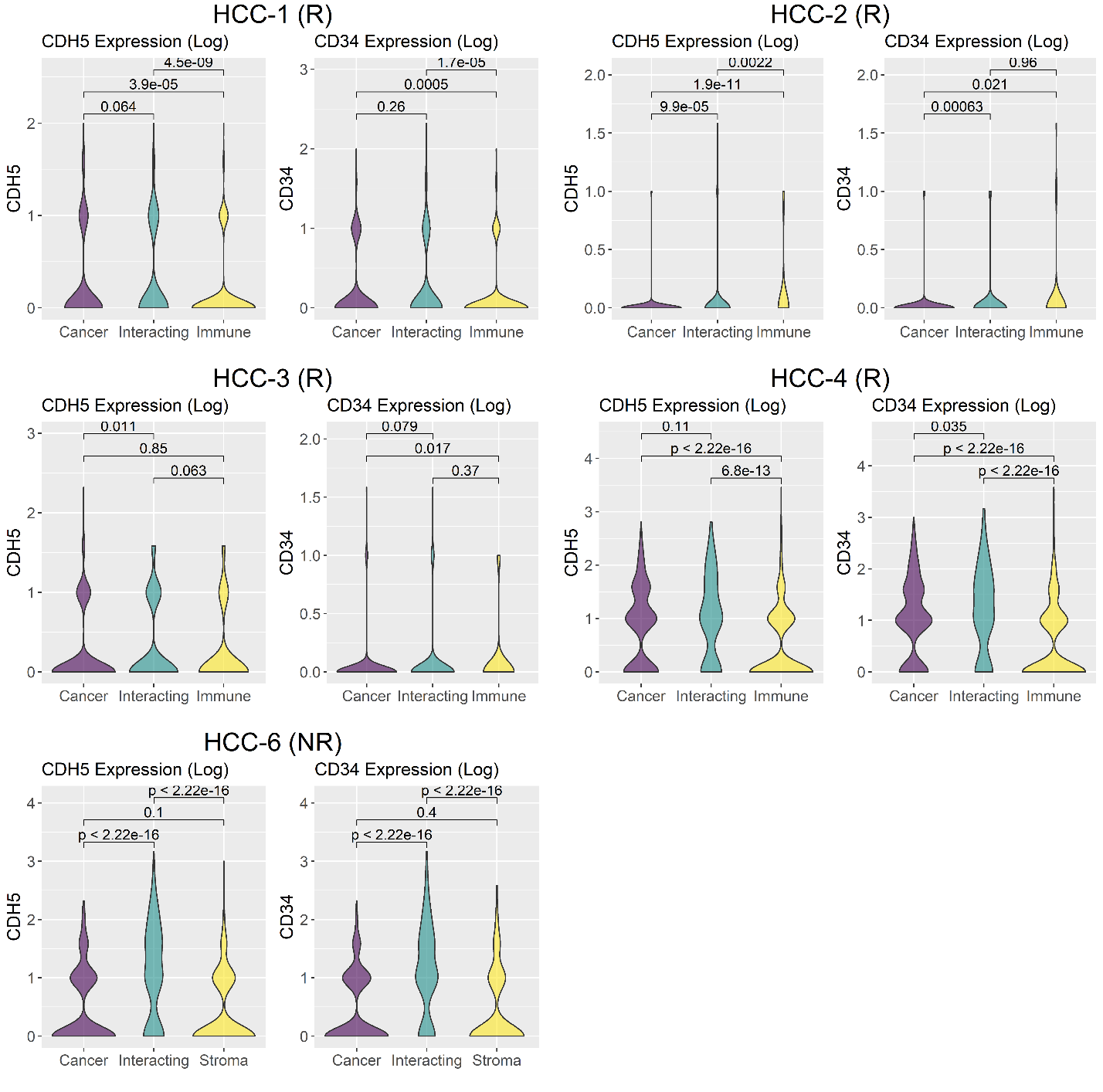
 Extended Data Fig. 10, **Vasculature marker expressions** Additional expression profiles of endothelial cell markers (*CDH5, CD34*) in spatial transcriptomic samples of 5 patients (responder: n=4, non-responder
